# Filtering Variables for Supervised Sparse Network Analysis

**DOI:** 10.1101/2020.03.12.985077

**Authors:** Lorin M. Towle-Miller, Jeffrey C. Miecznikowski, Fan Zhang, David L. Tritchler

## Abstract

**Motivation:** We present a method for dimension reduction designed to filter variables or features such as genes considered to be irrelevant for a downstream analysis designed to detect supervised gene networks in sparse settings. This approach can improve interpret-ability for a variety of analysis methods. We present a method to filter genes and transcripts prior to network analysis. This method has applications in a setting where the downstream analysis may include sparse canonical correlation analysis.

**Results:** Filtering methods specifically for cluster and network analysis are introduced and compared by simulating modular networks with known statistical properties. Our proposed method performs favorably eliminating irrelevant features but maintaining important biological signal under a variety of different signal settings. We show that the speed and accuracy of methods such as sparse canonical correlation are increased after filtering, thus greatly improving the scalability of these approaches.

**Availability:** Code for performing the gene filtering algorithm described in this manuscript may be accessed through the geneFiltering R package available on Github at https://github.com/lorinmil/geneFiltering. Functions are available to filter genes and perform simulations of a network system. For access to the data used in this manuscript, contact corresponding author.

**Contact:**, and

## 1 Background

As the costs of high-throughput experiments continue to decrease [13], it is common to assay a variety of genomic information from large cohorts of patients at the level of single nucleotide variants (SNVs) [5], gene expression level via ribonucleotide acid sequencing (RNA-Seq) [10] and, perhaps, at the metabolomic and proteomic levels [6]. In this setting, many sets of data are obtained where each set of variables is considered high-dimensional as it is obtained in a high-throughput setting. Analysis often proceeds with a combination of pathway and network analysis and it is believed that a better understanding in systems biology can be better detected through these integrated approaches of incorporating multiple ‘omics information together and observing the relationships between all datasets and the outcome of interest [12]. Conceptually, pathways and networks are similar but with certain distinctions as outlined in [3]. In short, pathways are small-scale systems while networks comprise genome- or proteome-wide interactions derived from integrative analyses of multiple datasets. Put another way, pathways are viewed as the “within” dataset collection of probes, while networks are the “between” datasets collection of probes where “probe” is a general term that could represent a single nucleotide polymorphism (SNP), a gene, a transcript, metabolite etc.

It is believed that certain complex diseases or clinical outcomes may be better predicted or understood through networks. For example, [4] notes a particularly interesting network in aggressive glioblastoma (GBM) where the network is discovered using the integration of copy number variation, gene expression, and gene mutation data obtained from the The Cancer Genome Atlas (TCGA) project [17]. TCGA project has supported the genomic data collection on approximately 11,000 patients across about 30 different types of tumors, where numerous data types were obtained through RNA-seq, MicroRNA sequencing, DNA sequencing, SNP detection, DNA methylation sequencing, and protein expression [17]. Most data types collected from TCGA studies and many other similar studies contain a large amount of features per data type for thousands of subjects. Network analyses that extend beyond just one type of data significantly raise the complexity of the analysis due to the high dimensionality and it introduces limitations due to computational concerns [2].

A challenge with data of this magnitude involves the computationally expensive modeling of the biological networks that can consume massive amounts of central processing unit (CPU) time, particularly for resampling based procedures such as bagging, boosting, bootstrapping or permutation which are used in various methods such as the penalized canonical correlation analysis (PCCA) method proposed in [20], the sparse canonical correlation analysis (SCCA) method proposed in [24], and the supervised penalized canonical correlation analysis method proposed in [16]. As expected, this problem gets worse with progressively larger patient cohorts as technological improvements allow for deeper assaying of genomic information. Many researchers performing network analyses currently have to find creative ways to remove genes due to memory constraints, such as removing genes with unknown location followed by averaging values across adjacent genes as done in [24] or removing features with low variance as done in [16]. The current use of dimension reduction techniques to computationally run many network analyses presents the need for filtering out noise prior to performing any primary analysis.

Our novel approach addresses this weakness by providing a scalable dimension reduction technique for filtering or removing genes/probes that are considered to be irrelevant from the customary downstream supervised network analysis approaches. An advantage of this approach is the improvement in speed of the network analysis algorithms such as SCCA. Another advantage is clarity of interpretation. That is, the biological understanding and interpretation of a network will be more easily discerned if the analysis does not include irrelevant or unimportant genes.

This work is based on extending a network filtering technique initially proposed in [18] where filters are made by applying k-means clustering of correlation estimates. Due to the generalizability of that network filtering calculation, it is straightforward and reasonable to extend it to a supervised network analysis setting where there are sets of genomic data on a cohort of subjects with an observed outcome (e.g. response to therapy) for those subjects.

### 1.1 Existing Filtering Methods

Few dimension reduction techniques have been proposed for pathway and network analyses. NARROMI, a noise reduction technique introduced in [26], utilizes mutual information to filter out unrelated features and is similar to the method described in this manuscript, but contains a limitation on handling only a single data type.

Principal component analysis (PCA) is another common method used to reduce dimensions for analyzing genes [14]. Performing PCA on data will combine similar features of the data into groups and distinguish the groups such that it encompasses the highest variance by utilizing linear algebra techniques. One major issue with PCA for network analysis is that it may remove too many features such that the sparsity assumption is unreasonable. Additionally, PCA is performed in an unsupervised setting with only one data type, making it less applicable to the network analysis framework.

Other similar methods, such as multidimensional scaling (MDS), are commonly used for reducing dimensionality while still maintaining the relationships. Enhancements of MDS, such as proposed in [19], could potentially maintain the sparsity assumption, however, this unsupervised approach would not consider the relationships with the outcome of interest. Additionally, it can only consider one data type at a time and ignores relationships across data types.

In Section 2 we introduce the proposed algorithm, in Section 3 we evaluate the performance of the algorithm under various simulations, and in Section 4 we apply the algorithm to a real world example. Our algorithm may be accessed freely in an R package with details provided in the abstract.

## 2 The Method

Our method applies to network analyses that include two data types and result in a continuous outcome. It is expected that a biological network will have probes within one data type that are related to features within another data type, which together result in the outcome. For example, suppose a small set of methylation sites lead to a small set of gene expression patterns, which ultimately result in an outcome for a particular disease. This network may be thought of as a casual chain with the outcome of interest, where the DNA methylation is the first data type and gene expression is the second intermediary data type. Furthermore, features in a given data type that are not included in the network should be jointly independent of the other data type features and the outcome.

With these network assuptions in mind, our proposed method utilizes estimated similarity matrices (e.g. Pearson correlation) between the outcome and the data types. A clustering method (e.g. k-means) is then applied to the similarities, and filters are established based on the clusters with the smallest means from the clustering results. This essentially removes any features in each data type that are not related to the other data type or the outcome. The full description of the algorithm is described in detail in Tables 1 and 2, and it is summarized in Figure 1.

**Figure 1:**
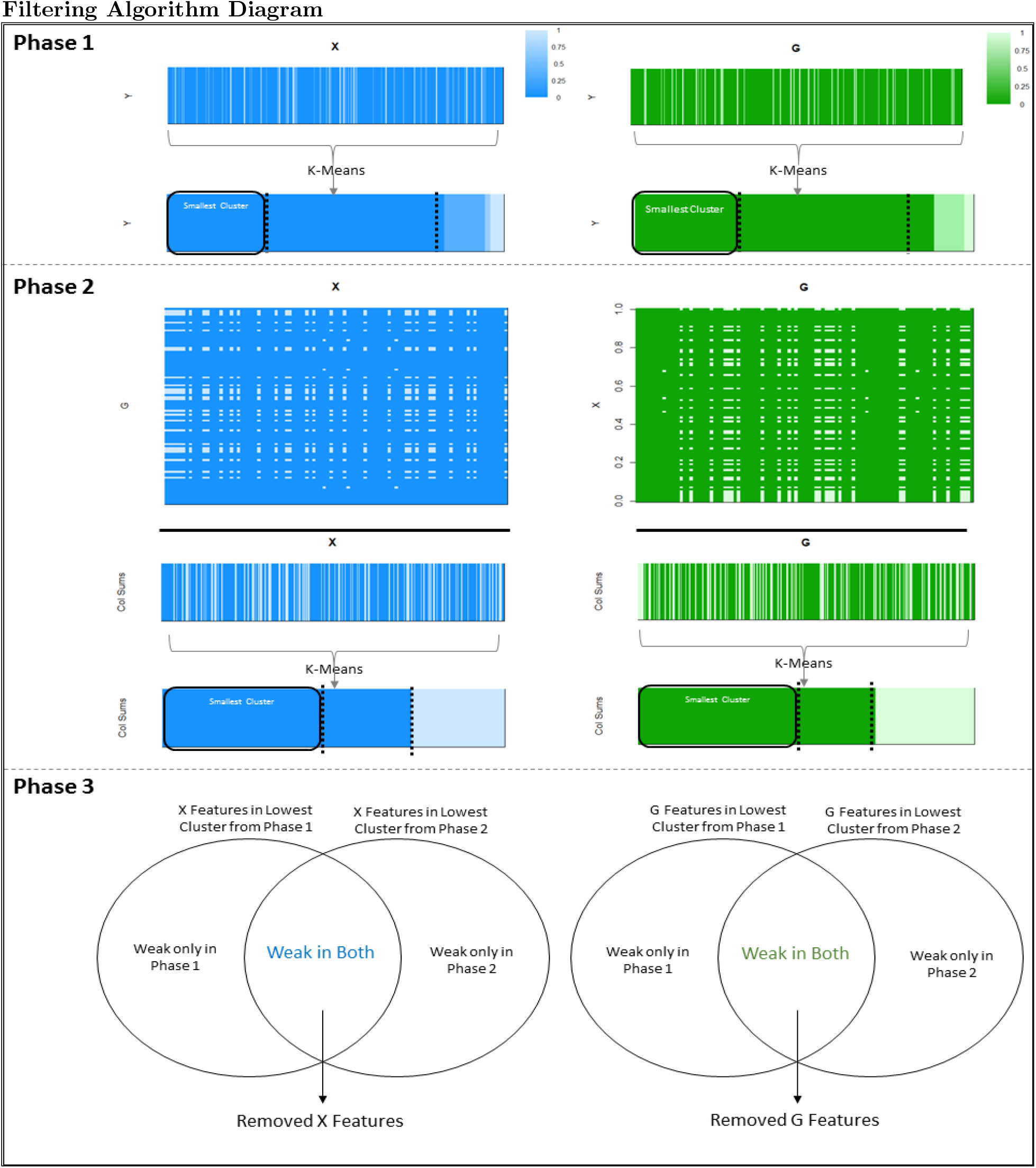
The filtering algorithm is performed in three phases. The phases are described in detail in Section 2 and Tables 1 and 2.

**Table 1:**
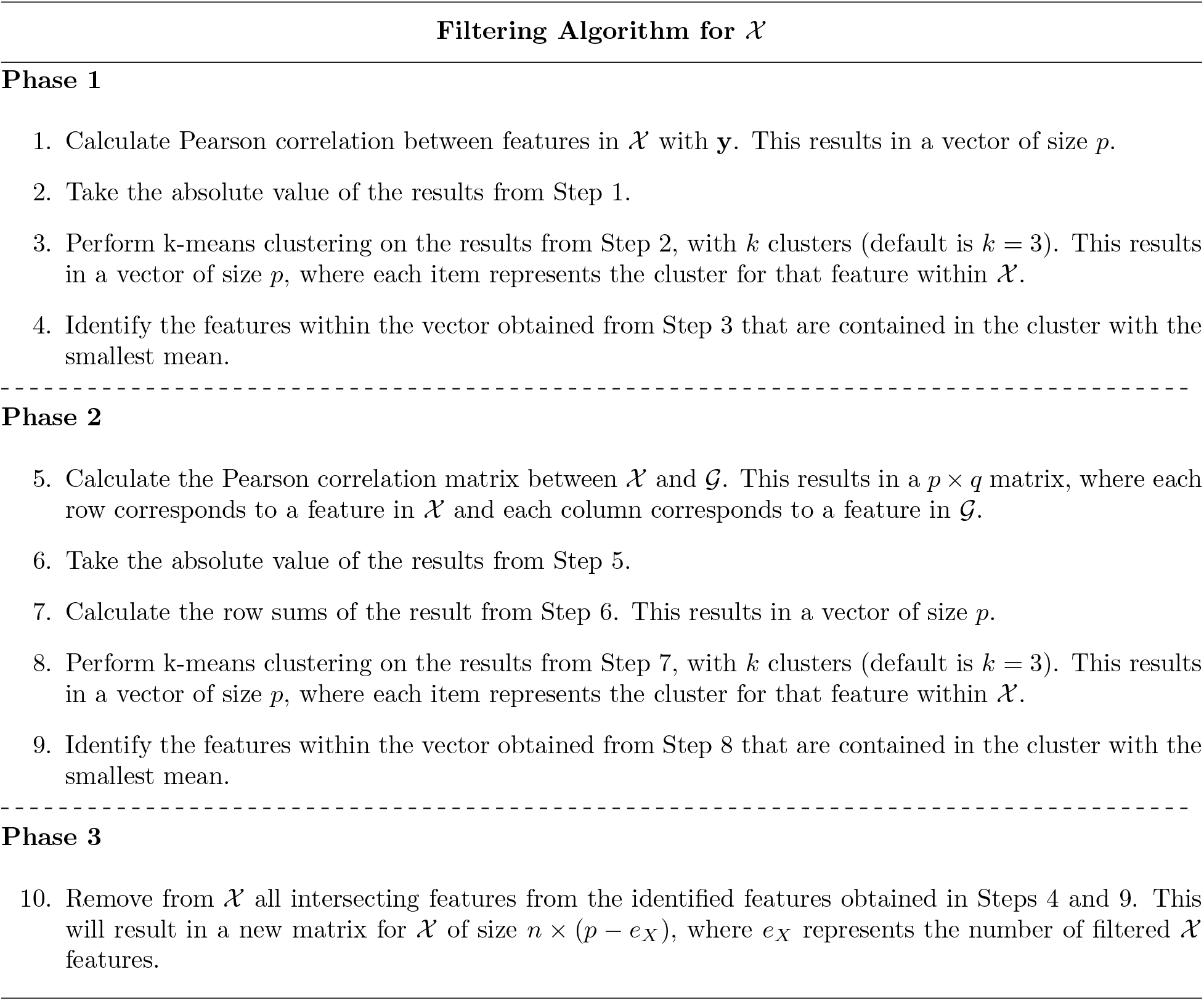
Filtering Algorithm for ***χ***. This algorithm is applied to ***χ*** prior to downstream analysis. The goal is to remove the features from ***χ*** that are neither correlated with **y** or ***ς*** The filters for ***χ*** are performed independently of the filters for ***ς***.

**Table 2:**
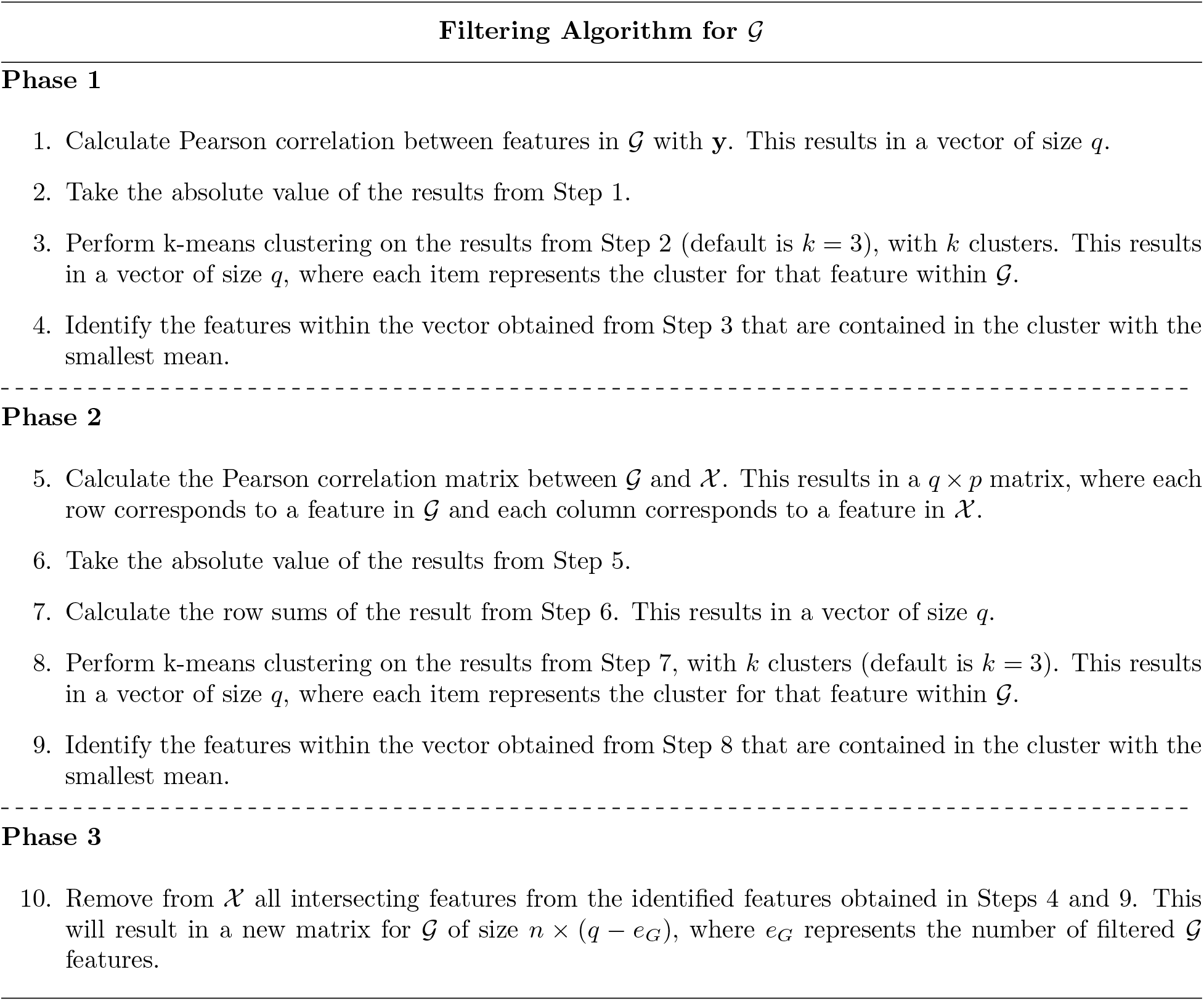
Filtering Algorithm for ***ς***. This algorithm is applied to ***ς*** prior to downstream analysis. The goal is to remove the features from ***ς*** that are neither correlated with **y** or ***χ***. The filters for ***ς*** are performed independently of the filters for ***χ***.

It should be emphasized that the goals of this algorithm are to remove irrelevant features within the data types prior to the downstream analysis and not to perform feature selection or identify the network features. Feature selection would aim at selecting features that are significantly related to the outcome and would likely eliminate too many “unneccesary” features, thus violating the sparsity assumption that is made in many network setting solutions. This filtering process is only intended for speeding up the downstream analysis and not to identify network features themselves. Figure 2 summarizes the intended workflow including this filtering algorithm for performing a network analysis.

**Figure 2:**
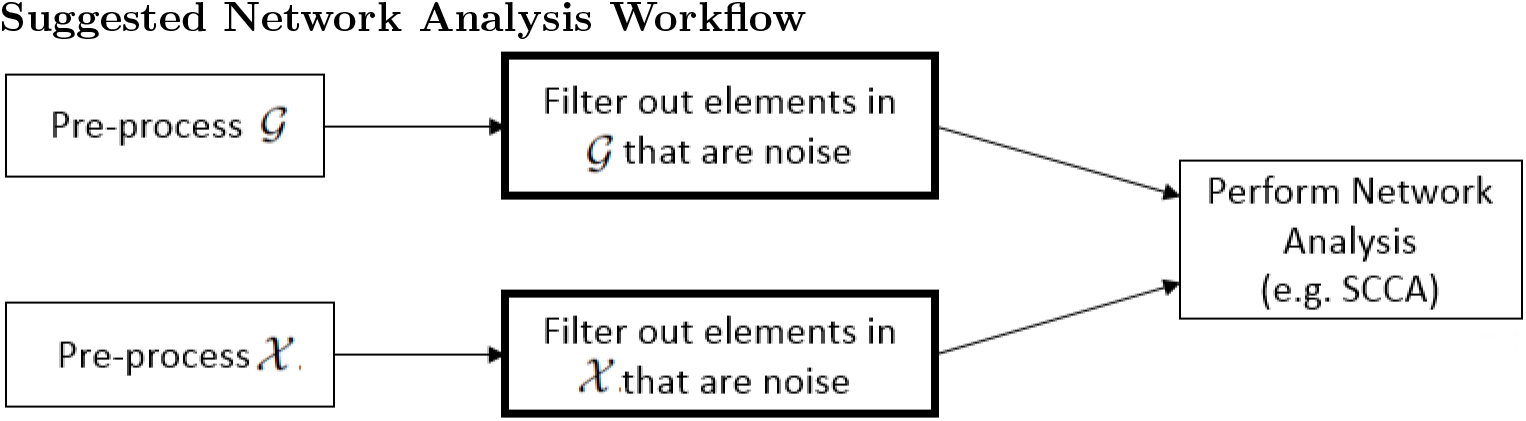
The proposed workflow when performing a network analysis with a filtering step between pre-processing (e.g. normalization and quality control measures) and the primary analysis in order to reduce run times for the time consuming network analysis methods.

### 2.1 Notation

Suppose there is an observed continuous outcome **y** for *n* subjects. Additionally, suppose that each subject has two omics data types collected, denoted as ***ς*** and ***χ***. There are *q* total features for each subject within ***ς*** and *p* total features for each subject within ***χ***, which results in ***ς*** and ***χ*** being *n × q* and *n × p,* respectively. Sub-matrices of sizes *n × q’* and *n × p’* are contained within ***ς*** and ***χ*** and make up the network features that explain the outcome, where there are *q’* network features in data type ***ς*** and *p*’ network features within data type ***χ***. In most network applications, a sparsity assumption is held, which would imply *q’ ≪ q* and *p’ ≪ p*.

### 2.2 Filtering Algorithm

Before the phases of the algorithm may be carried out, correlation matrices must first be estimated between ***ς*** and ***χ***, ***ς*** and **y**, and ***χ*** and **y**. Once all correlation matrices are estimated, the filtering method is then implemented.

The proposed filtering method comprises of three phases: Phase 1 gathers a list of features within both ***ς*** and ***χ*** to potentially be filtered based on correlations with **y**, Phase 2 gathers a list of features within both ***ς*** and ***χ*** to potentially be filtered based on correlations between ***ς*** and ***χ***, and Phase 3 combines the results from Phases 1 and 2 to establish a final set of features to be filtered. Phases 1 and 2 are performed separately, and Phase 3 will simply collect the intersection of results to construct a final set of features to be filtered. Note that filters on ***ς*** and ***χ*** are performed similarly and conducted independently, not conjointly. This will allow parallelization between the two data types in obtaining their set of features to be filtered.

In Phase 1, the absolute value of estimated correlations between **y** and all features within ***ς*** will be captured (resulting in a vector of size *q*), and k-means clustering will be applied to the correlation vector. All ***ς*** features that are contained within the cluster with the smallest mean will be marked for potential filtration. By grouping the features according to correlation strength with the outcome, a set of ***ς*** features that have the smallest correlations with the outcome may be identified. The appropriate number of clusters should be selected such that the cluster with the smallest mean adequately represents a set of features having low correlation with the outcome (suggesting that it would likely not be detected as a network feature in the subsequent network analysis). Additionally, the appropriate number of clusters should be small enough such that it selects a sufficient number of features in order to make a significant reduction on the analysis run times. A similar process would be repeated on all features within ***χ*** to identify a set of features under this data type that are also weakly correlated with the outcome **y**.

Phase 1 flags features for removal that are not related to the outcome, but does not address the correlation between data types. Hence, the goal of Phase 2 is to identify features within ***ς*** that are not correlated with features within ***χ***, and similarly to detect features within ***χ*** that do not appear to be correlated with features within ***ς***. The estimated correlation matrix between ***ς*** and ***χ*** is obtained and the absolute value of all estimated correlations is computed. For each feature of ***ς*** within this absolute correlation matrix, we sum across all features of ***χ*** (resulting in a vector of size *q*), and perform k-means clustering on the vector of summed correlations. All features of ***ς*** that are contained within the cluster with the smallest mean should be flagged for filtering as those features do not show strong correlation with features of ***χ***. A similar process for ***χ*** is conducted where the correlations across all ***ς*** features are summed and k-means clusters applied such that the features of ***χ*** contained in the cluster with the smallest mean are flagged for potential removal. Like Phase 1, the number of clusters should be selected so that the set of features captured adequately represent features that are weakly correlated with the corresponding data type while still flagging enough features such that it could make an impact on reducing run times for the primary analysis.

Phase 3 combines Phases 1 and 2 by selecting the set of features in ***ς*** and ***χ*** that were identified for filtering in both phases. Thus, by taking this intersection and removing it, we are removing features that are weakly correlated with **y** and weakly correlated within the corresponding data types. By definition of the networks of interest defined in Section 3, it is expected that the network features are correlated with the outcome and correlated to the other data type. Any features that violate both assumptions are likely features that will be removed.

## 3 Simulations

Simulations were performed to verify the validity and consistency of the filtering algorithm. The goals for the filtering algorithm are to retain all network features and maintain the sparsity assumption required for downstream methods while still removing enough features that would result in significant run time reductions.

We note many of the downstream network discovery techniques such as SCCA are based on sparse solutions where many of the variables or features are assigned zero coefficients. These sparse approaches are reasonable as it is believed only a relatively small set of features are involved in these functional network approaches. Thus, we need to ensure our filtering algorithm does not remove too many features such that a sparse solution is no longer appropriate. One criteria to assure a sparse setting is to examine the measure *n/log(p)* as described in [21] and ensure that *n/log(p)* is much larger than the total number of features within the active network(s), where n is the number of subjects and p is the total number of features within the data type. Since the total number of features within a network are not known in practical analyses, this measure is unobtainable for real data. However, for our simulations, the sparsity assumption is met when there is little change in the sparsity measure *n/log(p)* for both ***ς*** and ***χ*** when comparing before and after filtering.

For each simulation, we simulate matrices ***ς*** and ***χ*** with an outcome vector **y**. ***ς*** and ***χ*** are designed to contain network features which ultimately explain **y**. In addition to the network features, there may be other noise-like features within ***ς*** and ***χ***. For example, there may be features in ***ς*** and ***χ*** that are individually related to **y** but not part of a network while there may be features in ***ς*** and ***χ*** that are correlated with each other but not with **y**. Additionally there may be features that are purely noise uncorrelated with the outcome and uncorrelated with all other features. Ultimately, Figure 3 shows the types of features and correlation relationships that are contained in ***ς*** and ***χ***. As there are numerous types of relationships indicated in Figure 3 there will be numerous parameters to set in each simulation. In short, the model in Figure 3 is based on a multivariate normal setting and we note full details in the Appendix.

**Figure 3:**
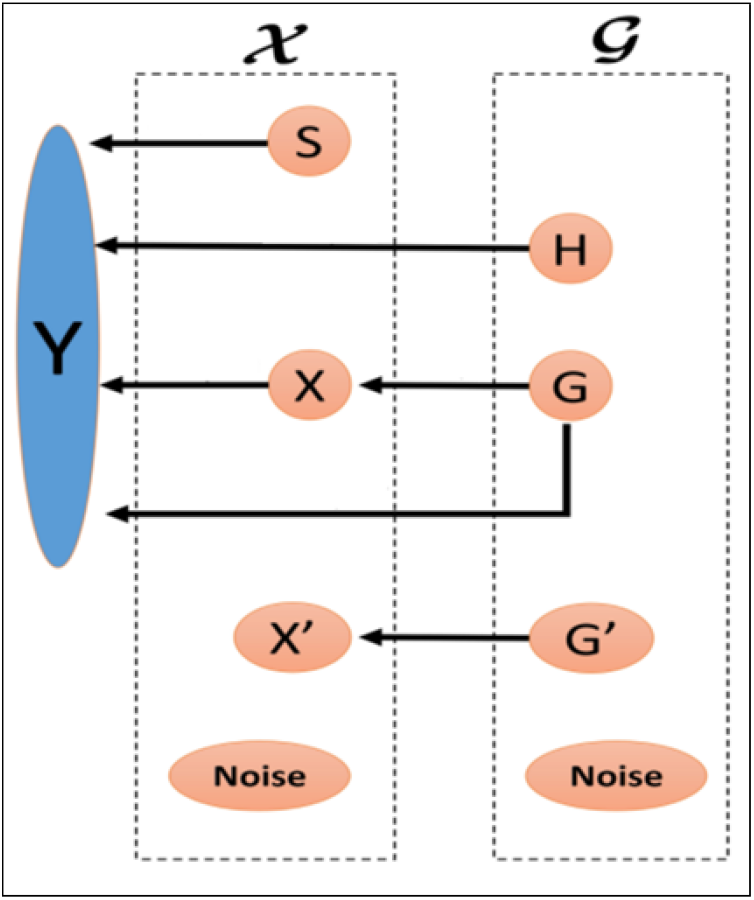
Diagram of latent variables in a network system with the outcome **y**. Each of the latent variables (**S**, **H**, **X**, **G**, **G**’, **H**’, and **Noise**) describe a subset of features within data types ***ς*** and ***χ*** with their corresponding relationships. **S** describes the set of features within ***χ*** that are related to **y** but not any features in ***ς***; **H** describes the set of features within ***ς*** that are related to **y** but not any features in ***χ***; **X’** describes the network features within ***χ*** which is related to the network features within ***ς*** and **y** (note that there are *p’* features in this subset); **G** describes the network features within ***ς*** which is related to the network features within ***χ*** and **y** (note that there are *q’* features in this subset); **X**’ describes the set of features within ***χ*** that are related to a set of features within ***ς*** but not related to **y**; **G**’ describes the set of features within ***ς*** that are related to a set of features within ***χ*** but not related to **y**; and **Noise** describes the features within ***χ*** and ***ς*** that are not related to each other or **y**. The variables within the respective dotted boxes summarize the various components contained within each of the data types, and the arrows signify a relationship and imply correlation between the corresponding components. Note that this figure was adapted from [25].

### 3.1 Parameter Selection

The parameters used in the simulations were adjusted to consider various levels of signal strength and various quantites of features. Parameters were chosen and assessed for weak, moderate, and strong network signal strength with the same number of subjects. Signal strengths were generated for a small, medium, and large number of total features (approximately 5000, 10000, and 20000 features within each data type, respectively). The number of features within the networks were kept small in order to accomodate the sparsity assumption and selected to reflect similar numbers described in [2]. Simulations were repeated 500 times under each setting to confirm reproducibility, and the filtering algorithm with *k* = 3 clusters under each phase was applied and assessed under each simulation separately. Refer to Table 3 for the details on the total number of features used for the simulations.

**Table 3:** The number of features across the simulations for the small, medium, and large number of features are summarized in the table. The same parameters for weak, moderate, and strong signal strengths were generated across each number of features for a total of 9 (3×3) different simulations, where each simulation contained 500 replications. The relationships between each of the components within ***χ*** and ***ς*** are described in Figure 3.

| Summary of Simulations |  |  |  |
| --- | --- | --- | --- |
| Item | Small | Medium | Large |
| Subjects | 1000 | 1000 | 1000 |
| Features in $\mathcal{X}$ | 5165 | 15165 | 25165 |
| Features in $\mathbf{S}$ | 50 | 50 | 50 |
| Features in $\mathbf{X}$ | 15 | 15 | 15 |
| Features in $\mathbf{X}'$ | 100 | 100 | 100 |
| Noise Features in $\mathcal{X}$ | 5000 | 15000 | 25000 |
| Features in $\mathcal{G}$ | 5140 | 10140 | 20140 |
| Features in $\mathbf{H}$ | 30 | 30 | 30 |
| Features in $\mathbf{G}$ | 10 | 10 | 10 |
| Features in $\mathbf{G}'$ | 100 | 100 | 100 |
| Noise Features in $\mathcal{G}$ | 5000 | 10000 | 20000 |

### 3.2 Results

The filtering algorithm described in Section 2 was applied to each of the simulations under all parameter settings. For each simulation, we examined the change in *n/log(p)* before and after filtering and the sensitivity of what was filtered. The reduction in computation time for executing SCCA on the data before and after filtering was also assessed on a subset of the simulations.

All simulation settings removed between 10 and 37 percent of features in both ***χ*** and ***ς*** with an average of 5 percent change in *n/log(p),* as summarized in Figure 4. Since the changes in this sparsity measure were minimal, it is suggested that the sparsity assumption has been maintained after filtration.

**Figure 4:**
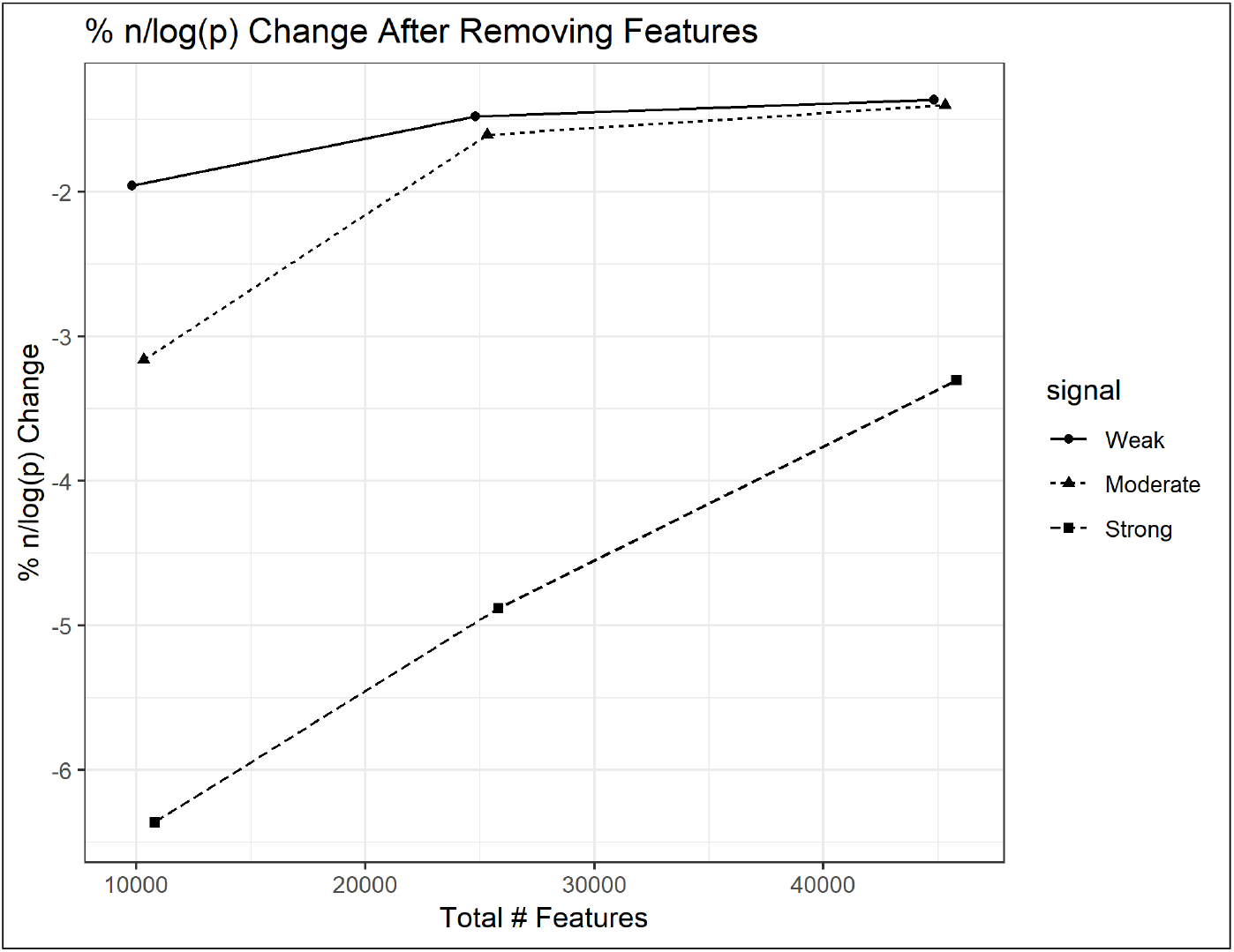
The average percent change in *n/log(p)* before and after removing features across the simulations from the nine different settings and the corresponding confidence interval bars. The points on each line correspond to small, medium, and large simulations from left to right. Also note that the total number of features is the number of features within both ***χ*** and ***ς*** combined, prior to any removing of features.

All simulations with strong signal did not remove any network features, and the weak and moderate signal strength simulations erroniously removed less than one network feature on average from both ***χ*** and ***ς***. Figure 5 displays representative simulations under the various signal strengths with a large number of features.

**Figure 5:**
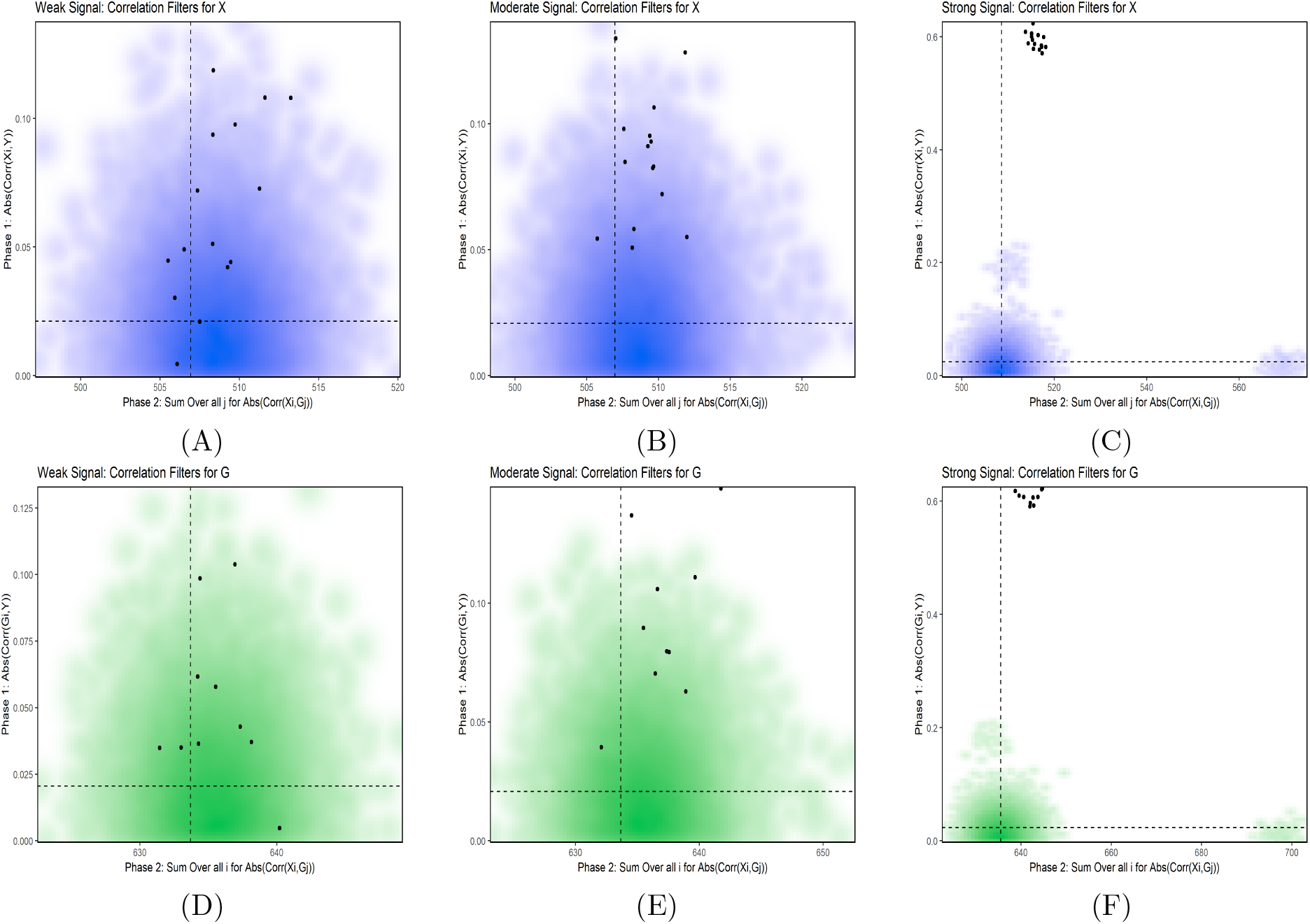
This figure shows representative simulations for weak, moderate, and strong signal strength under the simulations with a large number of features (45305). The black points represent the features that belong to the network. The dashed lines within each plot represents the k-means cluster threshold, where all points within the bottom left quadrant of each plot represent the features that will be removed. Plots (A) and (D) correspond to the correlation measures for ***χ*** and ***ς***, respectively, for a representative simulation under the weak signal setting; plots (B) and (E) correspond to the correlation measures for ***χ*** and ***ς***, respectively, for a representative simulation under the moderate signal setting; plots (C) and (F) correspond to the correlation measures for ***χ*** and ***ς***, respectively, for a representative simulation under the strong signal setting. In (A), we see 1 point (network feature) in the bottom left quadrant that would be erroneously removed.

To assess the reduction in run times for downstream analysis due to the filters in the number of features, the SCCA method described in [9] was employed on 64-bit Windows 10 with Intel Core i7-7700HQ, CPU of 2.80GHz, and 8.00 GB of RAM. Run times were collected under select simulations. The percent of run time reduction is summarized in Figure 6. The stronger the signal strength, the more features were removed which resulted in higher runtime reduction. For the simulations with a large number of features, the average run time for SCCA was 85.9 hours for the unfiltered data and 38.1 hours on the filtered data. For the simulations with a medium number of features, the average run time for SCCA was 4.4 hours for the unfiltered data and 2.1 hours on the filtered data. For the simulations with a small number of features, the average run time for SCCA was 35 minutes for the unfiltered data and 20 minutes on the filtered data. It is apparent that as the number of features increases, the amount of time to execute SCCA is exponentially increased. However, the steps proposed in this manuscript suggest that the filtering method may reduce run times for the primary analysis by up to about 80 percent.

**Figure 6:**
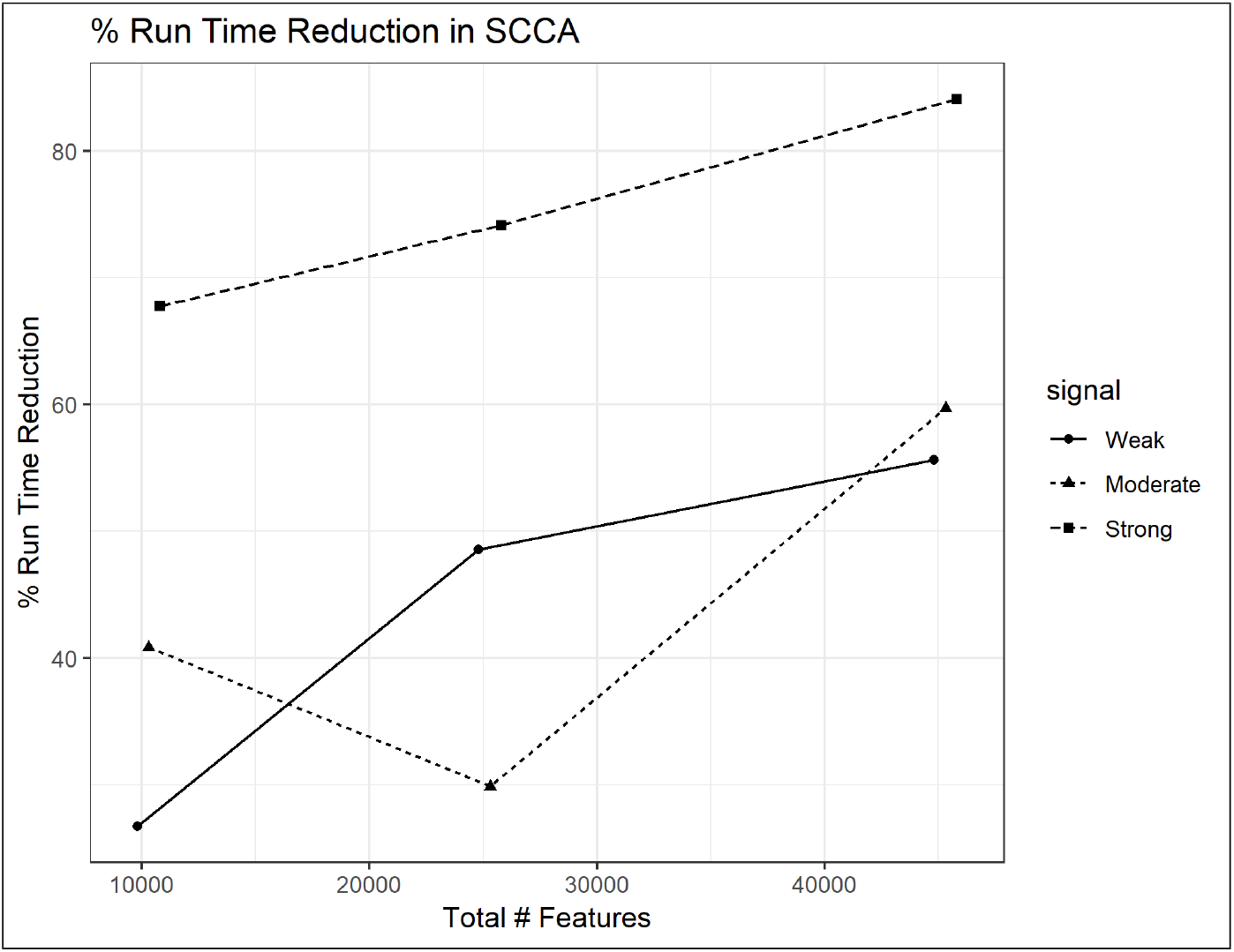
The average percent change in run times across the simulations from the different settings. The points on each line correspond to small, medium, and large simulations from left to right. Also note that the total number of features is the number of features within both ***χ*** and ***ς*** combined, prior to any filtering.

## 4 Application

According to [7], Uterine Corpus Endometrial Carcinoma (UCEC) is the fourth most common cancer amongst women in the United States with about 50,000 new cases reported and approximately 8000 deaths in 2013 alone. Due to the extensive prior research of UCEC data and sufficiently large obtainable sample size, UCEC was selected as one of the tumor types assessed in the Pan-Cancer analysis project within the TCGA database where mutation, copy number, gene expression, DNA methylation, MicroRNA, and RPPA were collected from the tumor samples [22].

**Table 4:** The results are averaged across 500 simulations under each setting. The goal is to remove enough features from ***ς*** and ***χ*** such that it makes an impact on computation time for the primary analysis but does not remove any network features, or effect the *n/log(p)* sparsity measure. There were some simulations with weak signal where a feature from the network was removed. Due to the weak signal and indirect relationship with **y**, it is not suprising that filters would erroneously remove features in these weak settings. It is also possible that these weakly related network features would not have been detecting in a downstream analyses such as SCCA.

| Summary of Simulation Filtering Results |  |  |  |  |
| --- | --- | --- | --- | --- |
| Signal Strength | Total # Features | Data Type | Features Removed | Network Features Removed |
| Weak | 45305 | $\mathcal{X}$ | 3376 | 0.34 |
| Weak | 45305 | $\mathcal{G}$ | 2704 | 0.22 |
| Moderate | 45305 | $\mathcal{X}$ | 3515 | 0 |
| Moderate | 45305 | $\mathcal{G}$ | 2684 | 0 |
| Strong | 45305 | $\mathcal{X}$ | 7339 | 0 |
| Strong | 45305 | $\mathcal{G}$ | 6364 | 0 |
| Weak | 25305 | $\mathcal{X}$ | 2089 | 0.34 |
| Weak | 25305 | $\mathcal{G}$ | 1389 | 0.21 |
| Moderate | 25305 | $\mathcal{X}$ | 2245 | 0.01 |
| Moderate | 25305 | $\mathcal{G}$ | 1507 | 0.01 |
| Strong | 25305 | $\mathcal{X}$ | 5791 | 0 |
| Strong | 25305 | $\mathcal{G}$ | 3729 | 0 |
| Weak | 15305 | $\mathcal{X}$ | 841 | 0.40 |
| Weak | 15305 | $\mathcal{G}$ | 835 | 0.15 |
| Moderate | 15305 | $\mathcal{X}$ | 1278 | 0.02 |
| Moderate | 15305 | $\mathcal{G}$ | 1263 | 0.01 |
| Strong | 15305 | $\mathcal{X}$ | 2242 | 0 |
| Strong | 15305 | $\mathcal{G}$ | 2132 | 0 |

It has been shown from prior studies that the p53 pathway is associated with many cancers, where p53 can be indirectly inactivated when MDM2 is overexpressed, PTEN or INK4A/ARF is mutated, the Akt pathway is deregulated, and other events occur [15]. We focus on chromosome 10, as it contains the PTEN gene and will utilize DNA methylation as one data type and gene expression as the other data type. Furthermore, prior research from [1] has shown that myometrial invasion is associated with tumor grade and patient survival, thus percent tumor invasion will be used as the outcome for this application.

The data collected within TCGA for UCEC contain 587 subjects with 485577 DNA methylation features and 56830 gene expression features. After focusing on chromosome 10, the final datasets resulted in 18901 methylation features, 2116 expression features, and 458 subjects.

The filtering method described in Section 2 applied with k = 3 clusters for each phase to the methylation and expression UCEC data removed 4106 methylation features and 333 expression features. The sparse canonical correlation analysis (SCCA) method described in [9] will be applied to the UCEC data both before and after the filtered datasets. The run time reduced from 48 minutes to 26 minutes after the filter is applied, and the same features selected from the analysis on the full set were also selected in the analysis from the filtered set.

In summary, consistency in SCCA results are maintained in this example while accomplishing run time reduction by almost 50 percent. Additionally, the percentage of features removed is consistent with the percentages found from the simulations, indicating our simulations are realistic with the size and scope of real data.

## 5 Discussion

There are several extensions of our method for consideration in future work. Our algorithm is currently limited to two types and is currently not dynamic to handle additional data types. In a setting with more than two data types, one approach to consider would be to calculate correlation matrices between each of the data types and apply filters where the relationships are expected. This approach would become more complex as the number of data types increases and would require prior network relationship knowledge. We also note the default version of our method has 3 clusters for all k-means calculations which works well in our simulations and application. However, in an extended setting with more data types, more research may be needed to tune this parameter such that it is removing enough features to make an impact on run time reduction while not removing too many features leading to a violation on the sparsity assumption.

Another limitation in our method is that it assumes that the networks are approximately linearly correlated. If the network had a complex relationship that was not linear, this would limit the applicability of the filtering method. However, many common downstream analyses such as CCA based methods also assume that the networks are linearly related. We also caution that our approach should not be combined with downstream testing based approaches as our supervised approach will introduce a bias that will violate downstream multiple testing corrections designed to control a Type I like error.

Our method also does not model for additional subject covariates that may be available for analysis. One ad hoc approach in this setting may be to compute residuals from a model involving the outcome and additional covariates. These residuals can then serve as the “outcome” in our filtering method. We are exploring this approach in a future manuscript.

We note that the estimate of run time reduction in Figure 6 is based on Monte Carlo simulations of modest size and not the entire 500 simulations as employed for the other simulation estimates. This is due to the abundant amount of resources required per deployment of the SCCA procedure. The limited resources for running SCCA under these settings further validate the need for reducing the data. In addition, our method was applied and tested in R version 3.4.4 [11]. Due to memory constraints in R, it cannot allocate extremely large matrices. This poses a problem when calculating correlation matrices where the number of features within ***ς*** and ***χ*** are large. To accommodate these constraints, our method is applied on a row by row basis which significantly slows down the filtering method run times. Although the filtering method would only need to be applied once, other techniques and programming languages may be considered in obtaining the correlation matrices in order to avoid the memory limitations in R.

## 6 Conclusions

Network analysis techniques such as in [8], [25] and [23] are critical for the understanding of large collections of datasets such as TCGA project. However, they are computationally intensive and as such would greatly benefit from dimension reduction such as gene filtering. We note our filtering approach improves computational times and biological interpretability while also maintaining a sparse setting required for most downstream penalization based network discovery algorithms. By reducing run times, more resources can be spent on the formal analysis while still maintaining consistent results and assumptions.

## 7 Author Contributions

LT-M assisted in algorithm conceptualization, performing simulations and real data application, and drafted the manuscript. JM assisted in algorithm conceptualization and drafting manuscript. DT assisted in algorithm conceptualization. FZ assisted with coding of simulations, real data application, and SCCA implementation.

## 8 Acknowledgements

Research reported in this publication was partially supported by the National Center for Advancing Translational Sciences of the National Institutes of Health under award Number UL1TR001412. The content is solely the responsibility of the authors and does not necessarily represent the official views of the NIH.

## Notes

### Competing Interest Statement

The authors have declared no competing interest.

### Summary of Updates

Statement removed to declare submission.

## References

[1] R. Boronow, C. Morrow, W. Creasman, P. Disaia, S. Silverberg, A. Miller, and J. Blessing. Surgical Staging in Endometrial Cancer: Clinical-pathologic Findings of a Prospective Study. Obstetrics and Gynecology, 63(6):825–832, 1984.

[2] L. Chin, W. C. Hahn, G. Getz, and M. Meyerson. Making Sense of Cancer Genomic Data. Genes & Development, 25(6):534–555, 2011.

[3] P. Creixell, J. Reimand, S. Haider, G. Wu, T. Shibata, M. Vazquez, V. Mustonen, A. Gonzalez-Perez, J. Pearson, C. Sander, et al. Pathway and Network Analysis of Cancer Genomes. Nature Methods, 12(7):615, 2015.

[4] C. Danussi, U. D. Akavia, F. Niola, A. Jovic, A. Lasorella, D. Pe’er, and A. Iavarone. RHPN2 Drives Mesenchymal Transformation in Malignant Glioma by Triggering RhoA Activation. Cancer Research, 2013.

[5] X. Dong, L. Zhang, B. Milholland, M. Lee, A. Y. Maslov, T. Wang, and J. Vijg. Accurate Identification of Single-Nucleotide Variants in Whole-Genome-Amplified Single Cells. Nature Methods, 14(5):491, 2017.

[6] M. Larance and A. I. Lamond. Multidimensional Proteomics for Cell Biology. Nature Reviews Molecular Cell Biology, 16(5):269, 2015.

[7] D. A. Levine, C. G. A. R. Network, et al. Integrated Genomic Characterization of Endometrial Carcinoma. Nature, 497(7447):67, 2013.

[8] J. C. Miecznikowski, D. P. Gaile, X. Chen, and D. L. Tritchler. Identification of Consistent Functional Genetic Modules. Statistical Applications in Genetics and Molecular Biology, 15(1):1–18, 2016.

[9] E. Parkhomenko, D. Tritchler, and J. Beyene. Sparse Canonical Correlation Analysis with Application to Genomic Data Integration. Statistical Applications in Genetics and Molecular Biology, 8(1):1–34, 2009.

[10] M. Pertea, D. Kim, G. M. Pertea, J. T. Leek, and S. L. Salzberg. Transcript-Level Expression Analysis of RNA-seq Experiments with HISAT, StringTie and Ballgown. Nature Protocols, 11(9):1650, 2016.

[11] R Core Team. R: A Language and Environment for Statistical Computing. R Foundation for Statistical Computing, Vienna, Austria, 2018.

[12] V. K. Ramanan, L. Shen, J. H. Moore, and A. J. Saykin. Pathway Analysis of Genomic Data: Concepts, Methods, and Prospects for Future Development. TRENDS’ in Genetics, 28(7):323–332, 2012.

[13] J. A. Reuter, D. V. Spacek, and M. P. Snyder. High-throughput Sequencing Technologies. Molecular Cell, 58(4):586–597, 2015.

[14] M. Ringnér. What is Principal Component Analysis? Nature Biotechnology, 26(3):303, 2008.

[15] S. Surget, M. P. Khoury, and J.-C. Bourdon. Uncovering the Role of p53 Splice Variants in Human Malignancy: A Clinical Perspective. OncoTargets and Therapy, 7:57, 2014.

[16] A. Thum, S. Mönchgesang, L. Westphal, T. Lubken, S. Rosahl, S. Neumann, and S. Posch. Supervised Penalized Canonical Correlation Analysis. 05 2014.

[17] K. Tomczak, P. Czerwinska, and M. Wiznerowicz. The Cancer Genome Atlas (TCGA): An Immeasurable Source of Knowledge. Contemporary Oncology, 19(1A):A68, 2015.

[18] D. Tritchler, E. Parkhomenko, and J. Beyene. Filtering Genes for Cluster and Network Analysis. BMC Bioinformatics, 10(1):193, 2009.

[19] J. Tzeng, H. H.-S. Lu, and W.-H. Li. Multidimensional Scaling for Large Genomic Data Sets. BMC Bioinformatics, 9(1):179, 2008.

[20] S. Waaijenborg, P. C. V. de Witt Hamer, and A. H. Zwinderman. Quantifying the Association Between Gene Expressions and DNA-markers by Penalized Canonical Correlation Analysis. Statistical Applications in Genetics and Molecular Biology, 7(1), 2008.

[21] M. J. Wainwright. Sharp Thresholds for High-Dimensional and Noisy Sparsity Recovery Using L1 Constrained Quadratic Programming Lasso. IEEE Transactions on Information Theory, 55(5):2183–2202, 2009.

[22] J. N. Weinstein, E. A. Collisson, G. B. Mills, K. R. M. Shaw, B. A. Ozenberger, K. Ellrott, I. Shmulevich, C. Sander, J. M. Stuart, C. G. A. R. Network, et al. The Cancer Genome Atlas Pan-cancer Analysis Project. Nature Genetics, 45(10):1113, 2013.

[23] D. M. Witten, R. Tibshirani, and T. Hastie. A Penalized Matrix Decomposition, with Applications to Sparse Principal Components and Canonical Correlation Analysis. Biostatistics, 10(3):515–534, 2009.

[24] D. M. Witten and R. J. Tibshirani. Extensions of Sparse Canonical Correlation Analysis with Applications to Genomic Data. Statistical Applications in Genetics and Molecular Biology, 8(1):1–27, 2009.

[25] F. Zhang, J. Miecznikowski, and D. Tritchler. Identification of Supervised and Sparse Functional Genomic Pathways. SUNY University at Buffalo, Department of Biostatistics, Technical Report, (1801), 2018.

[26] X. Zhang, K. Liu, Z.-P. Liu, B. Duval, J.-M. Richer, X.-M. Zhao, J.-K. Hao, and L. Chen. NARROMI: A Noise and Redundancy Reduction Technique Improves Accuracy of Gene Regulatory Network Inference. Bioinformatics, 29(1):106–113, 2012.

